# Identification of CP77 as the third orthopoxvirus SAMD9L inhibitor with a unique specificity for a rodent SAMD9L

**DOI:** 10.1101/551556

**Authors:** Fushun Zhang, Xiangzhi Meng, Michael B Townsend, Panayampalli Subbian Satheshkumar, Yan Xiang

**Author notes:** To whom correspondence should be addressed: Yan Xiang. Contribute equally (Fushun Zhang, Xiangzhi Meng).

## Abstract

Orthopoxviruses (OPXVs) have a broad host range in mammalian cells, but Chinese hamster ovary (CHO) cells are non-permissive for vaccinia virus (VACV). Here, we revealed a species-specific difference in host restriction factor SAMD9L as the cause for the restriction and identified orthopoxvirus CP77 as a unique inhibitor capable of antagonizing Chinese hamster SAMD9L (chSAMD9L). Two known VACV inhibitors of SAMD9 and SAMD9L (SAMD9&L), K1 and C7, can bind human and mouse SAMD9&L, but neither can bind chSAMD9L. CRISPR-Cas9 knockout of chSAMD9L from CHO cells removed the restriction for VACV, while ectopic expression of chSAMD9L imposed the restriction for VACV in a human cell line, demonstrating that chSAMD9L is a potent restriction factor for VACV. Contrary to K1 and C7, cowpox virus CP77 can bind chSAMD9L and rescue VACV replication in cells expressing chSAMD9L, indicating that CP77 is yet another SAMD9L inhibitor but has a unique specificity for chSAMD9L. Binding studies showed that the N-terminal 382 amino acids of CP77 were sufficient for binding chSAMD9L and that both K1 and CP77 target a common internal region of SAMD9L. Growth studies with nearly all OPXV species showed that the ability of OPXVs in antagonizing chSAMD9L correlates with CP77 gene status and that a functional CP77 ortholog was maintained in many OPXVs, including monkeypox virus. Our data suggest that species-specific difference in rodent SAMD9L poses a barrier for cross-species OPXV infection and that OPXVs have evolved three SAMD9L inhibitors with different specificities to overcome this barrier.

**IMPORTANCE:** Several OPXV species, including monkeypox virus and cowpox virus, cause zoonotic infection in humans. They are believed to use wild rodents as the reservoir or intermediate hosts, but the host or viral factors that are important for OPXV host range in rodents are unknown. Here, we showed that the abortive replication of several OPXV species in a Chinese hamster cell line was caused by a species-specific difference in the host antiviral factor SAMD9L, indicating that SAMD9L divergence in different rodent species poses a barrier for cross-species OPXV infection. While the Chinese hamster SAMD9L could not be inhibited by two previously identified OPXV inhibitors of human and mouse SAMD9L, it can be inhibited by cowpox virus CP77, indicating that OPXVs encode three SAMD9L inhibitors with different specificity. Our data suggest that OPXV host range in broad rodent species depends on three SAMD9L inhibitors with different specificities.

## INTRODUCTION

The orthopoxvirus (OPXV) genus of the poxvirus family consists of more than ten closely-related species, including the infamous human pathogen variola virus (VARV, causative agent for smallpox), the smallpox vaccine vaccinia virus (VACV), and emerging zoonotic agents monkeypox virus (MPXV) and cowpox virus (CPXV) (1). These viruses as well as ectromelia virus (ECTV), camelpox virus (CMLV) and teterapox virus (TATV) are known as the Old World OPXVs for originating from the Eurasian continent, while raccoonpox virus (RNCV), skunkpox virus (SKPV) and volepox virus (VPXV) are recognized as the North American OPXVs for being endemic in North America (2). VARV has been eradicated from nature, but the cessation of smallpox vaccination and the waning of anti-OPXV herd immunity have increased the risk of zoonotic OPXV infections. MPXV is highly virulent in humans (3), and the recent increase in human monkeypox cases across a wide geographic area is a concern for global health security (4). CPXV and VACV are responsible for zoonoses in Europe, Asia and South America (5, 6). Novel OPXV species have also been discovered in recent human cases (7, 8), including Akhmeta virus (AKMV) isolated from the town of Akhmeta, Georgia (country).

OPXVs vary greatly in their animal host range (2), with some, such as VARV, CMLV and TATV, only known to infect a single mammalian species, while others, such as MPXV, CPXV and VACV, capable of infecting a wide variety of mammalian hosts. OPXVs are often named after the host in which they were first isolated, while their reservoir hosts in nature are unknown. However, OPXV infections are often associated with contact with rodents. CPXV is carried and believed to be transmitted to humans and domestic animals by bank voles and striped field mice in Western Europe (9, 10). MPXV is likely carried by rope squirrels in Africa (11). The spread of VACV in Brazil involves wild rodents as the reservoir or intermediate hosts (12). Contrary to their varied host range in nature, OPXV host range in tissue culture cells are almost universally broad (13). It was thus noteworthy that the Chinese hamster ovary (CHO) cells were found to be nonpermissive for VACV, owing to a rapid shutoff of protein synthesis that resulted in a block of viral intermediate mRNA translation (14, 15). A CPXV gene from the Brighton Red (BR) strain, CP77, could rescue VACV replication in CHO cells (16). CPXV BR CP77 encodes a 668 amino acid (aa), 77-kDa protein with nine predicted ankyrin repeats and a C-terminal F-box domain (17). CP77 is conserved in all CPXV strains, having greater than 91% aa sequence identity amongst different strains. A CP77 ortholog with approximately 90% aa identity is also present in many MPXV strains. In contrast, the CP77 ortholog is either deleted or fragmented in all sequenced VACV strains, while a large deletion occurs in VARV CP77 ortholog that results in a protein of at most 452 aa.

Although the specific host pathway that restricts VACV replication in CHO cells and the mechanism by which CP77 overcomes the restriction are unknown, CP77 is long recognized as a host-range gene that provides a similar function as the VACV host-range genes K1L and C7L (18, 19). K1L encodes a 284 aa protein consisted entirely of ankyrin repeats (20), while C7L encodes a 150 aa protein forming a single β-sandwich (21). VACV with deletions in both K1L and C7L (vK1^−^C7^−^) replicate abortively in many mammalian cell lines (19); the replication defect in human cells could be rescued by K1L, C7L or CP77. K1L and CP77 are specific to OPXVs, whereas functional C7L homologs are present in nearly all mammalian poxviruses (22). Recent studies revealed that K1 and C7 provide an equivalent function for viral growth in human and mouse cells by inhibiting a common set of host restriction factors, SAMD9 (23, 24) and SAMD9L (25). We thus investigated in this study whether CP77 functions similarly to K1 and C7 by directly targeting SAMD9 or SAMD9L. Our results not only identified CP77 as yet another OPXV inhibitor of SAMD9L but also revealed a species-specific difference in SAMD9L that explains the specific requirement of CP77 for OPXV tropism in some rodent cells.

## MATERIALS AND METHODS

### Cells and viruses

VERO (ATCC CCL-81), BSC40 (ATCC CRL-2761), BT20 (ATCC HTB-19), and CHO-K1 (ATCC CCL-61) were originally from ATCC. HEK 293FT was from Thermo Fisher Scientific (cat. no. R70007). WT VACV WR strain, K1L and C7L deletion VACV (vK1^−^C7^−^) and a panel of vK1^−^C7^−^-derived recombinant viruses expressing VACV-K1 (vVACV-K1L) or a C7 homolog from different poxviruses (vVACV-C7L, vYLDV-67R, vMYXV-M62R, vMYXV-M63R, vMYXV-M64R, vSPPV-063 and vSWPV-064) were described before (22, 26, 27). Other OPXVs (28), ECTV (Moscow), CMLV (CMS), MPXV (West African clade), AKMV (29), SKPV (USA1978-WA), VPXV (USA1985-CA), TATV (Dahomey1968) and RNCV (MD1964_85A) were from CDC and cultured in BSC-40 cells.

VACVs expressing WT or mutated CPXV CP77 were constructed as follows. An intermediate virus named vK1^−^C7^−^CP77^−^ was first constructed by deleting the CP77 gene fragments from vK1^−^C7^−^ (22) with the transient dominant selection method, as described previously (30). The deletion of the CP77 gene fragment in the recombinant virus was confirmed by PCR amplification of the CP77 region of the virus. Viruses expressing V5-tagged CP77 were derived from vK1^−^C7^−^CP77^−^ through homologous recombination with a transfer plasmid. The transfer plasmid contains (i) 500 bp of downstream flanking region of CP77, (ii) a GFP under the control of VACV late promoter P11, (iii) CPXV BR CP77 with a C-terminal V5 tag, and (iv) ∼500 bp of upstream flanking region of CP77 including the CP77 promoter. Specific mutations of CP77 were introduced into the plasmid through recombinant PCR as described previously (26). All constructs were confirmed by DNA sequencing. The recombinant virus construction was done according to standard protocols (31). In brief, the transfer plasmids were transfected into VERO cells that were infected with vK1^−^C7^−^CP77^−^. Recombinant viruses expressing GFP were picked under the fluorescence microscope and purified through four rounds of plaque isolation on VERO cells.

### SAMD9 and SAMD9L expression constructs

chSAMD9 ORF was PCR-amplified with the primer pair (5’-GGATGACGATGACAAGGCAGAGAAACTCAACCTTCCAGAGA-3’ and 5’-GGCTCCGCGGTTAGACAATTTTAATGTCATAAGCA-3’) from cDNA synthesized from CHO cellular mRNAs. A 3xFlag tag sequence was then appended to the 5’ end of the ORF by PCR, and the final PCR product was cloned between KpnI and SacII sites of the pcDNA3.1/V5-His-topo vector (Thermo Fisher Scientific). The chSAMD9 cDNA was completely sequenced and found to be identical to the Chinese hamster SAMD9 reference sequence in GenBank (XM_016963772.1). chSAMD9L ORF was cloned similarly but with the primer pair (5’-TGACGATGACAAGAATGAACAAGTAACTGCACCTAAATTGG-3’ and 5’-GGCTCCGCGGTTAGATTACTTTTATGCCATATGCCAGAGG-3’). The chSAMD9L cDNA was found to be identical to the Chinese hamster SAMD9L sequence XM_027389843.1 except for a difference at codon 618 (T to C substitution) that did not result in an amino acid change. Plasmids for expressing NTPase-TPR domains of SAMD9&L were constructed similarly but with PCR primers that only amplify the specific region of SAMD9&L.

### Generation of CHO cells with gene knockouts

The plasmids used for the gene knockout were constructed from lentiCRISPRv2 (Addgene plasmid #52961) according to the published protocol (32). In brief, lentiCRISPRv2 was digested with BsmBI and ligated with a pair of oligonucleotides with the specific guide sequence. For each target gene, two guide sequences were designed with the web tool CRISPOR (33). They were as follows: 5’-AGTCATTTGTATTCTCTGGA-3’ (chSAMD9 #1), 5’-CAAAAGAGGATGTGAATCTG-3’ (chSAMD9 #2), 5’-CAATGAAGAAGTGACAGGGA-3’ (chSAMD9L #1), 5’-ATCAGAAGTGCTGGACCCCG-3’ (chSAMD9L #2). The lentiCRISPRv2-derived plasmids and the packaging plasmids (pMD2.G and psPAX2) were transfected into HEK 293FT cells to produce lentiviruses, which were transduced into CHO cell as described (25). The transduced cells were subjected to puromycin (15 μg/ml) selection for 7 days. SAMD9 or SAMD9L genotype of the cells was identified by sequencing as described previously (25). The cellular genomic DNA was extracted using the QIAamp DNA blood mini kit (Qiagen). ∼500 bp of DNA flanking the target site was PCR-amplified, cloned to pGEM-T vector (Promega), and the sequence of 10-20 clones determined by Sanger sequencing. The primer pairs for chSAMD9 and chSAMD9L are (5’-AAGAGAGCTGGGGATAATGC-3’ and 5’-GATTTCTGCAGTTCCTTGAA -3’) and (5’-AAGTAATCATATGACTACATGTAA-3’ and 5’-GTGTTCTTTTATTGAGAGCT-3’).

### Generation of a cell line that expresses chSAMD9L under an inducible promoter

chSAMD9L was cloned into pCW57.1 (Addgene #41393), an “all-in-one” doxycycline inducible lentiviral vector with rtTA-VP16-2A-puro. In an effort to increase the expression level of the inducible gene, the lentiviral vector was modified by introducing F67S and R171K substitution into rtTA (34) and by replacing the minimal promoter downstream of the Tet Response Element with EF1 alpha promoter. The plasmid was used in making lentiviruses for transduction of BT20 cells as described above. Transduced cells were selected with puromycin at 3 μg/ml.

### Luciferase assay

Cells were infected with VACV WR that expressed a luciferase reporter under the control of either late p11 promoter (35) or the synthetic early/late (S E/L) promoter (36), in the presence or absence of cytosine arabinoside (AraC). Cells were lysed with buffer and luciferase activity measured at 8 or 24 hour post infection (hpi) according to the manufacturer’s instructions (Promega).

### Viral growth analysis

Cells in 12-well plates were incubated with 1 PFU per cell of different viruses for 2 h at room temperature. Following adsorption, the cells were washed twice with phosphate-buffered saline (PBS). One set of the cells was harvested immediately as the 0 hpi sample, while others were moved to an 37°C incubator to initiate viral entry and harvested at different times post infection. The viral titers in the cell lysates were determined by plaque assays on VERO cells.

### Immunoprecipitation and Western blot analysis

The binding of SAMD9&L with viral proteins was determined by co-immunoprecipitation as described before (25). In brief, 293FT cells were transfected with the SAMD9 or SAMD9L expression plasmid and then infected with different VACVs. The cells were lysed on ice with a lysis buffer (0.1% (w/v) NP-40, 50 mM Tris, pH 7.4, 150 mM NaCl), and the cleared cell lysates were mixed with V5-agarose beads (Sigma-Aldrich) for 30 min at 4°C. After washing with lysis buffer, the beads were resuspended in SDS sample buffer, the eluted proteins were resolved by SDS-PAGE and detected with Western blot as described previously (26). The detection antibodies were mouse monoclonal antibodies (mAb) against V5 (Sigma-Aldrich; clone V5-10) and Flag tag (Sigma-Aldrich).

## RESULTS

### Chinese hamster SAMD9 and SAMD9L have unique binding specificity for poxvirus proteins

All known poxvirus antagonists of SAMD9&L were able to bind their targets in mammalian cells (25, 37). To assess whether they could also inhibit SAMD9&L from Chinese hamster, we first set out to study their binding to Chinese hamster SAMD9&L. Chinese hamster SAMD9 (chSAMD9) and SAMD9L (chSAMD9L) were cloned from CHO cells into a mammalian expression vector and transiently expressed in HEK 293FT cells. The cells were then infected with a panel of vK1^−^C7^−^ derived VACV expressing different V5-tagged viral proteins, including one that expressed CPXV CP77 (vCPXV-CP77). The ability of the V5-tagged viral protein to bind the Flag-tagged SAMD9 or SAMD9L was assessed by immunoprecipitation (IP) with anti-V5 antibody followed by Western blot. All known poxvirus SAMD9&L antagonists, including K1, C7 and C7 homologs from diverse mammalian poxviruses, failed to precipitate chSAMD9L (Fig. 1A), in contrast to the binding of SAMD9&L from human and mouse by nearly all these viral proteins (25, 37). Interestingly, CP77 was able to precipitate chSAMD9L, suggesting that it is also a SAMD9L inhibitor and has a unique specificity for chSAMD9L. All the tested viral proteins except for K1 failed to precipitate chSAMD9 (Fig, 1B). Altogether, SAMD9&L from Chinese hamster stand out among the SAMD9&L that have been characterized so far for their resistance to binding by many of the known poxvirus SAMD9&L inhibitors.

**Figure 1.**
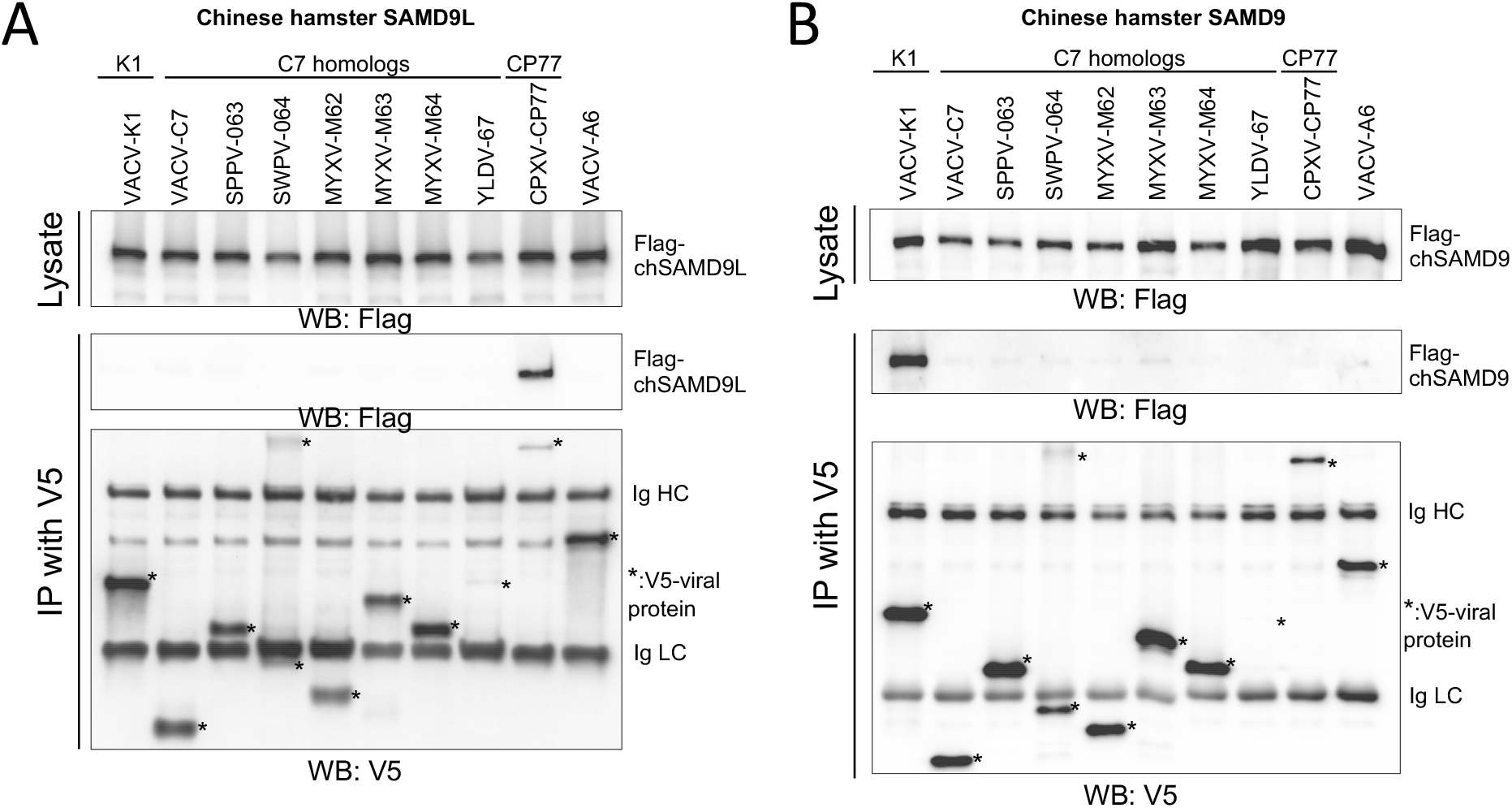
Only CPXV CP77 can bind Chinese hamster SAMD9L, while only VACV K1 can bind Chinese hamster SAMD9. 293FT cells were transfected with a plasmid expressing Flag-tagged chSAMD9L **(A)** or chSAMD9 **(B)** and infected with a panel of vK1^−^C7^−^-derived VACVs that expressed different V5-tagged viral proteins. VACV-A6, a viral protein involved in viral assembly, was used as a negative control. The cell lysates were immunoprecipitated with anti-V5 antibody. Epitope-tagged proteins in the cell lysate and precipitate were detected with anti-Flag or anti-V5 antibody in Western blot. The heavy and light chains of the precipitated antibody (Ig HC and LC) serve as loading controls. * indicates V5-tagged proteins.

### chSAMD9L is required for restricting VACV replication in CHO cells; CP77 is a SAMD9L inhibitor with a unique specificity for chSAMD9L

The lack of binding to chSAMD9 or chSAMD9L by K1 and C7 suggest that a failure of VACV in antagonizing chSAMD9 or/and chSAMD9L could be the reason why VACV replicates abortively in CHO cells. To test this idea, we knocked out either chSAMD9 or chSAMD9L from CHO cells with CRISPR-Cas9. For each gene knockout (KO), two independent KOs with different guide sequences (named as #1 and #2) were performed, and the pooled KO cells were tested for permissiveness for the panel of vK1^−^ C7^−^-derived VACVs. Similar to the parental CHO cells, the two chSAMD9 KO CHO cells (named ΔSAMD9) were nonpermissive for all recombinant VACVs except for vCPXV-CP77, which grew more than 100-fold in titer after 24 h of infection (Fig. 2A&B). In contrast, the two chSAMD9L KO cells (named ΔSAMD9L) were permissive for vK1^−^C7^−^ as well as all its derivatives (Fig. 2C&D). Several cell clones were also isolated and found to be similar to the pooled cells in permissiveness for VACV (Fig. 2E). The specific KO of either chSAMD9 or chSAMD9L in the cell clones were confirmed by sequencing the region targeted by CRISPR-Cas9 guide. Indels that result in frameshift were found (Fig. 2F). These data show that chSAMD9L but not chSAMD9 is required for restricting VACV replication in CHO cells and that CP77 has a unique capability of antagonizing chSAMD9L. The well-characterized ΔSAMD9L clone 2F was used in all subsequent experiments.

**Figure 2.**
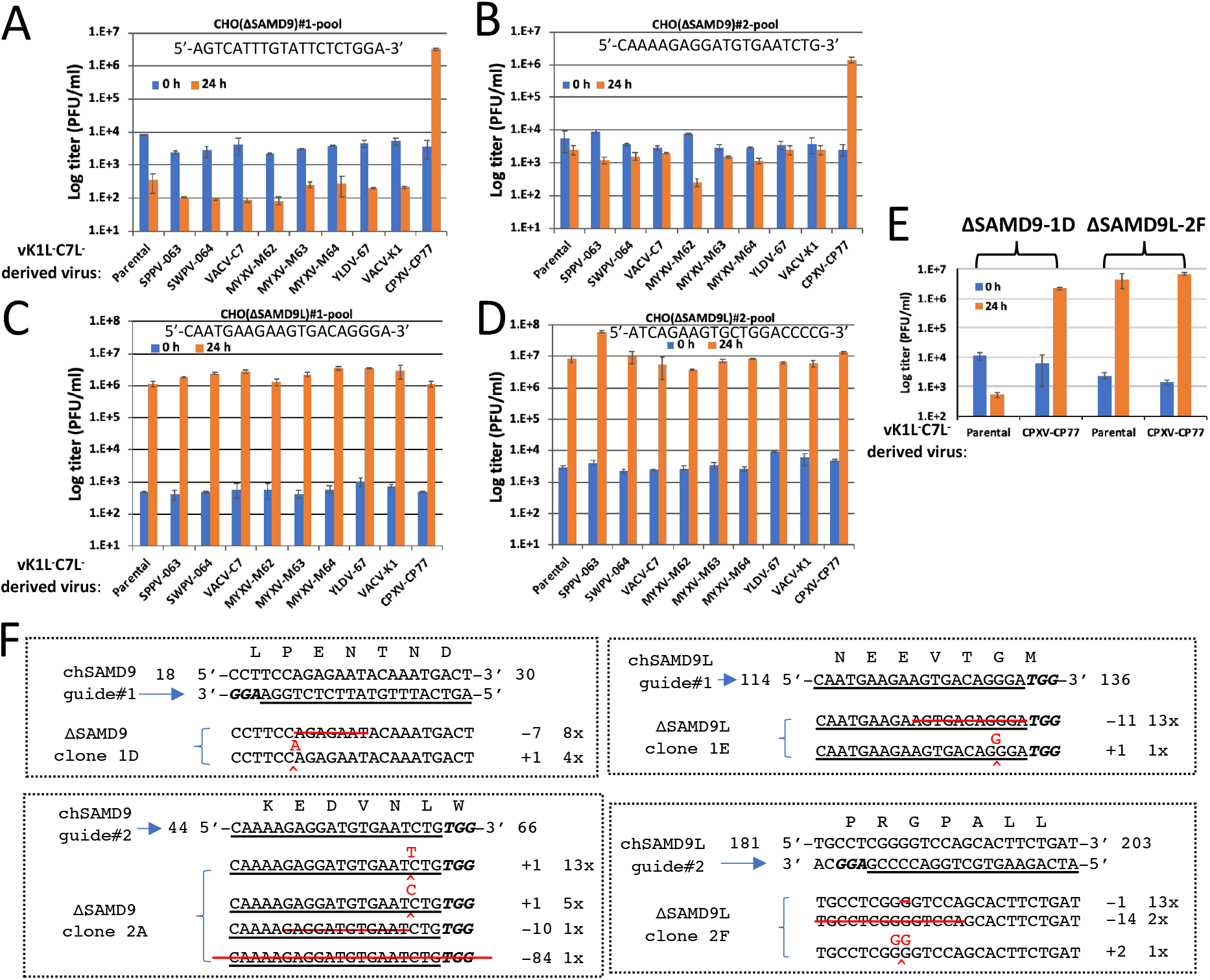
The restriction of VACV by CHO cells can be abolished by knocking out chSAMD9L. (A-D). CHO cells were transduced with lentiviral vectors expressing Cas9 and a guide sequence targeting either chSAMD9 (A&B) or chSAMD9L (C&D). For each gene, two independent KOs with different guide sequences (named as #1 and #2, shown in graph) were performed, and the pooled KO cells were infected with the panel of vK1^−^C7^−^-derived VACVs at a MOI of 1 PFU/cell. Viral titers at 0 and 24 hour post-infection (hpi) were measured by plaque assay in VERO cells. **(E).** Several KO cell clones were isolated from the pooled cells. Infection studies were performed on the cell clones and the results were similar to A-D. Rrepresentative data with ΔSAMD9-1D clone and ΔSAMD9L-2F clone are shown. **(F).** The genotype of the cell clones was determined by sequencing. The guide sequence is underlined with the PAM sequence in bold italics. Starting and ending positions of the guide in the ORF and the encoded amino acid sequence are also shown. Shown below the target are the genomic sequences from cell clones. The red line indicates deletion. ^ indicates insertion. The number after the + and -denotes the number of indels, and the number before the “x” denotes the number of times the sequence was detected from a total of 10-20 cloned PCR products.

It was previously shown that VACV replication in CHO cells was blocked at translation of post-replicative mRNA (18). To assess the effect of SAMD9L KO on viral early and post-replicative protein synthesis, we infected the parental and ΔSAMD9L CHO cells with VACVs that expressed luciferase gene under the control of either the synthetic early/late promoter or the late p11 promoter. In some infections, viral DNA replication was blocked to allow only early protein synthesis. The early luciferase expression measured at 8 hpi was at a comparable level in BSC40 cells and ΔSAMD9L CHO cells but was reduced by 10-fold in the parental CHO cells (Fig. 3A). A similar reduction on early luciferase expression was also observed at 24 hpi (Fig. 3B). However, the greatest difference between the parental and ΔSAMD9L CHO cells was observed on post-replicative, late luciferase expression, which was reduced by ∼10,000-fold in the parental CHO cells (Fig. 3B). Altogether, the data indicates that chSAMD9L KO alleviated the block on viral protein synthesis, particularly the late protein synthesis.

**Figure 3.**
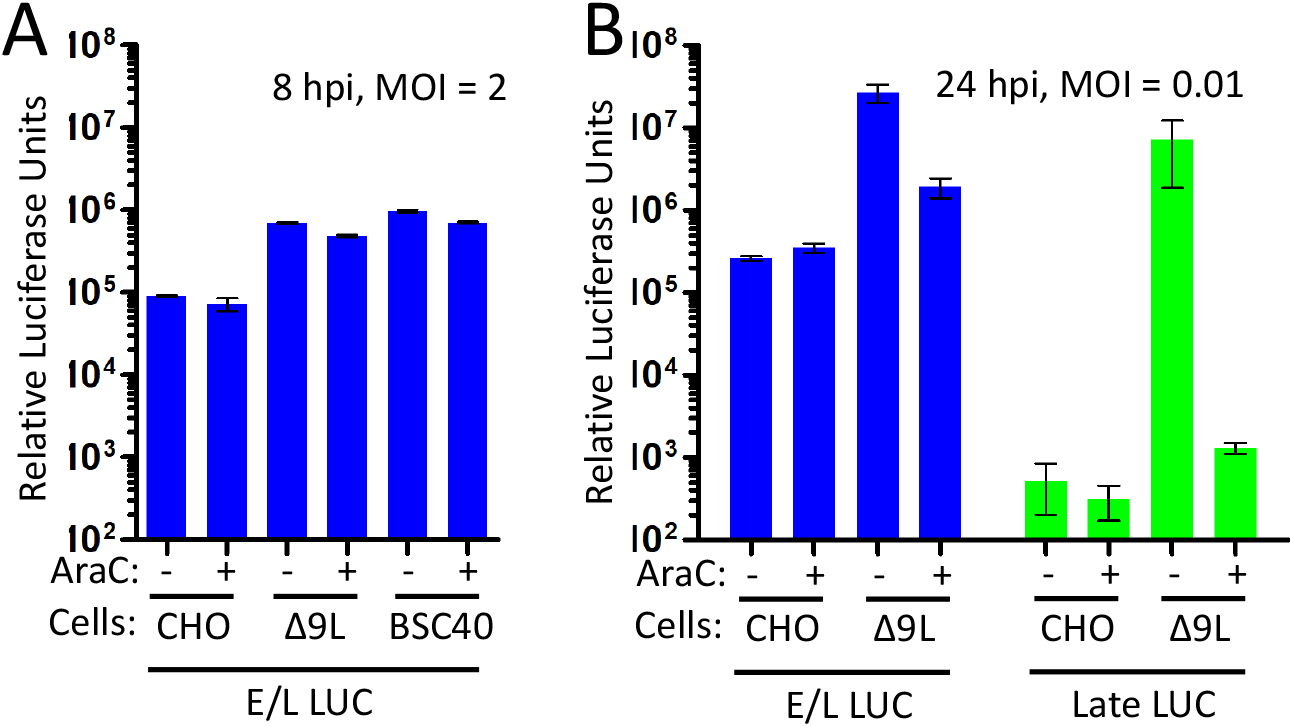
The block in viral protein synthesis in CHO cells can be abolished by knocking out chSAMD9L. BSC40 cells, the parental and ΔSAMD9L (2F clone) CHO cells were infected with VACVs that either expressed a luciferase reporter under the control of either the synthetic early/late (E/L) promoter or the late p11 promoter, in the presence or absence of AraC. The cells were lysed after either 8 or 24 h of infection and luciferase (LUC) activities were measured.

### Expression of chSAMD9L in human cells is sufficient for recapitulating poxvirus restriction property of CHO cells

While many mammalian cell lines were nonpermissive for vK1^−^C7^−^, we found some human cell lines, including the breast cancer BT20 cells, expressed a low level of human SAMD9 and were thus permissive for vK1^−^C7^−^. We made a stable BT20 cell line that expressed chSAMD9L by transduction with a lentivirus encoding chSAMD9L controlled by the tetracycline-dependent promoter. The cell line (named i-chSAMD9L) expressed chSAMD9L only when induced with doxycycline, in a dose-dependent manner (Fig. 4A). When not induced to express chSAMD9L, i-chSAMD9L cells were fully permissive for vK1^−^C7^−^ (or its derivatives), which grew by ∼1000-fold in titer after 24 h of infection (Fig. 4B). When induced to express chSAMD9L, the cell line became nonpermissive for vK1^−^C7^−^ and its derivatives that either expressed K1 (vVACV-K1) or C7 (vVACV-C7) (Fig. 4B), recapitulating CHO cells in terms of the restriction of poxvirus. Moreover, vCPXV-CP77 was able to grow ∼100-fold in titer after 24 h of infection, again demonstrating that CP77 can antagonize chSAMD9L.

**Figure 4.**
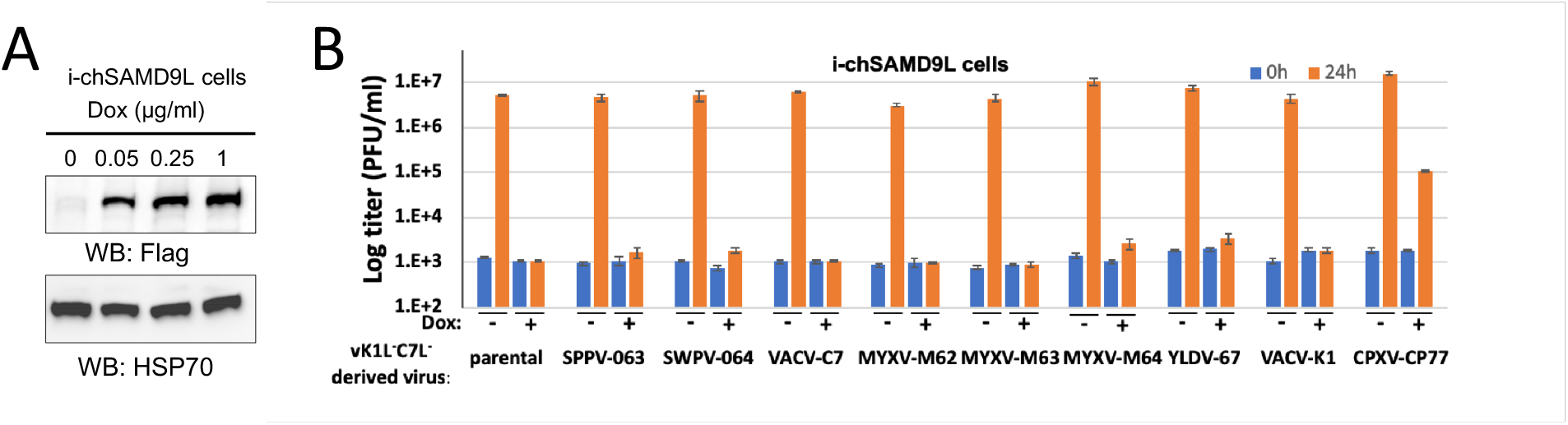
Expression of chSAMD9L in human cells is sufficient for recapitulating poxvirus restriction property of CHO cells. **(A).** A stable human breast cancer BT20 cell line with inducible expression of chSAMD9L was established via lentiviral transduction. The cell line (i-chSAMD9L) was cultured with medium containing the indicated concentration of doxycycline (Dox). The level of chSAMD9L and the control HSP70 protein in the cell lysates was determined by Western blot. (B). The i-chSAMD9L cells were either untreated or treated with 1 μg/ml Dox. The cells were then infected with the panel of vK1^−^C7^−^-derived VACV. Viral titers at 0 and 24 hpi were measured by plaque assay in VERO cells.

### K1 and CP77 target the same region of SAMD9&L

A computational analysis predicated SAMD9&L having the following domains from the N-to the C-terminus (38): SAM, AlbA, SIR2, P-loop NTPase, TPR, and OB (Fig. 5A). The N-terminal 385 aa of human SAMD9 (hSAMD9), which contains the predicted SAM and AlbA domains, was reported to be sufficient for binding to a poxvirus C7 homolog (39). To find out which region of hSAMD9 is targeted by K1, we constructed hSAMD9 mutants with deletions in different domains and tested the binding of the mutants to K1. We found aa 607-1172 of hSAMD9, which contains the putative NTPase and TPR domains, was sufficient for binding to K1 but not C7 (Fig. 5B). This binding was disrupted by two specific K1 substitution mutations (S2C#2 or S1-mut6) that were previously shown to disrupt K1 host-range function in human cells (20). Furthermore, both K1 and CP77, but not C7, can bind to the similar region of mouse SAMD9L (aa 598-1172), but only CP77 can bind to this region of chSAMD9L (aa 594-1172) (Fig. 5B). Thus, the binding to NTPase-TPR domain by K1 and CP77 displays the same species-specificity as to the full-length protein. Altogether, the data demonstrate that K1 and CP77 target the same NTPase-TPR region of SAMD9&L, differing from the region that is targeted by C7.

**Figure 5.**
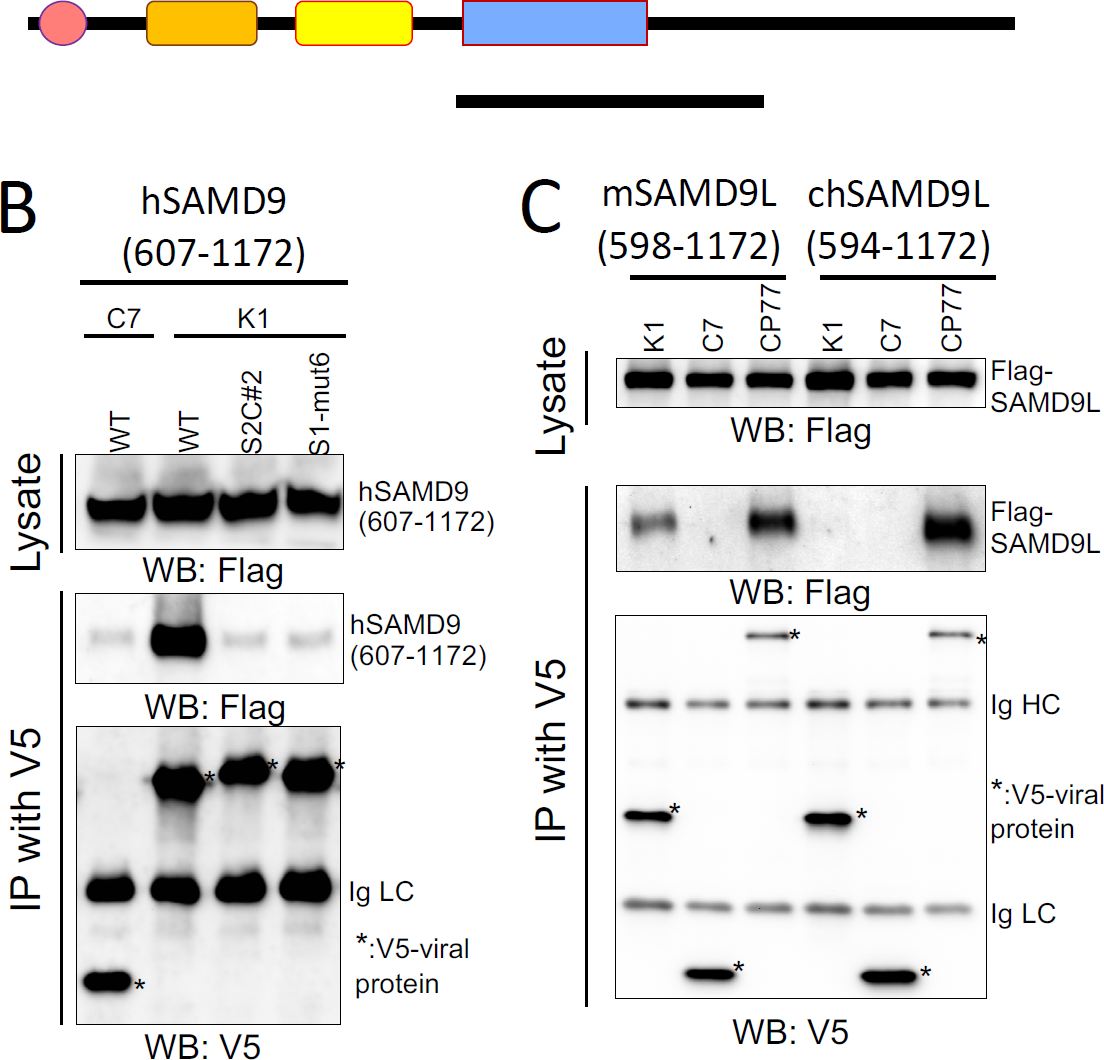
K1 and CP77 target a common internal region of SAMD9&L. **(A).** Schematics of the predicted SAMD9&L domain architecture and the SAMD9&L truncation with only the NTPase-TPR domain. **(B&C)**. 293FT cells were transfected with a plasmid expressing the putative NTPase and TPR domains of SAMD9&L (aa 607-1172 for hSAMD9, aa 598-1172 for mSAMD9L, and aa 594-1172 for chSAMD9L) and infected with VACV expressing C7, K1 or CP77. Co-IP and Western blot were performed as described in Fig. 1. The two K1 substitution mutations (S2C#2 or S1-mut6) were previously shown to disrupt K1 host-range function in human cells (20).

### The N-terminal 382 aa of CP77 is sufficient for binding to chSAMD9L

The full-length CP77 of 668 aa was predicted to contain 9 ankyrin repeats with a C-terminal F-box domain (Fig. 6A). Only the N-terminal 352 aa was reported to be necessary for the function of CP77 in CHO cells, and the deletion of ankyrin repeat 5 could disrupt CP77 function (40). To determine whether chSAMD9L binding has a similar requirement for CP77 residues, we constructed recombinant VACV from vK1^−^C7^−^ by inserting truncated CP77 into the viral genome. The recombinant virus expressing the N-terminal 382 aa of CP77 (ΔC, maintaining the N-terminal seven ankyrin repeats) can grow in CHO cells, although with ∼10-fold reduction in yield compared to the virus expressing the full-length CP77 (Fig. 6B). In contrast, the virus expressing a CP77 with a deletion in aa 235-266 (the predicted ankyrin repeat 5, Δ5) failed to grow in CHO cells. Correspondingly, CP77-ΔC can precipitate both the full-length and the NTPase-TPR region of chSAMD9L, whereas CP77-Δ5 cannot precipitate either (Fig. 6C). A reduced amount of full-length chSAMD9L was precipitated by CP77-Δc than by full-length CP77, correlating with the reduced growth in CHO cells by the CP77-Δc-expressing VACV. Compared to the full-length CP77, CP77-ΔC was also detected at a reduced level in the infected cells (Fig. 6C).

**Figure 6.**
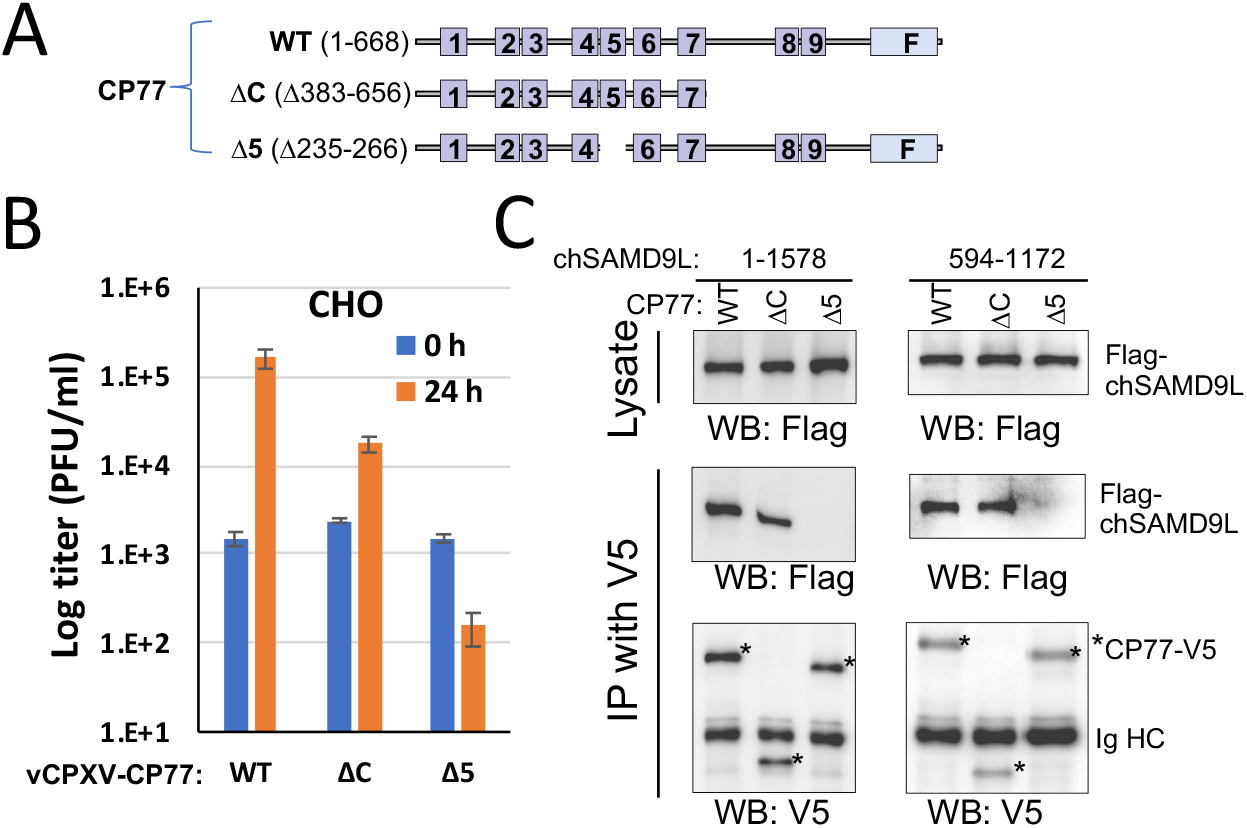
The N-terminal 382 aa of CP77 is sufficient for binding to chSAMD9L. (A). Schematics of different CP77 constructs. Ankyrin repeats are shown as boxes and numbered. The C-terminal F-box is indicated with “F”. ΔC: deletion of aa 383-656 (maintaining the N-terminal seven ankyrin repeats); Δ5: deletion of aa 235-266 (ankyrin repeat 5). (B). CHO cells were infected with vK1^−^C7^−^-derived VACV expressing either WT or mutated CP77. Viral growth was determined as described in Fig. 2. (C). Full-length and aa 594-1172 of chSAMD9L were subjected to co-immunoprecipitation with either WT or mutated CP77 as described in Fig. 1.

### The ability of OPXV in antagonizing chSAMD9L corresponds to their CP77 gene status

CP77 gene is variably maintained and diversified in different OPXV species (Table 1). To assess the ability of chSAMD9L in restricting different OPXV species, we compared the growth of all available OPXV species in the parental and ΔSAMD9L CHO cells (Fig. 7). ECTV and CMLV are similar to VACV for growing in ΔSAMD9L CHO cells but failing to grow in the parental CHO cells, indicating that chSAMD9L is also a restriction factor for ECTV and CMLV. The CMLV CP77 ortholog is largely deleted with only ∼155 nucleotides left, while the ECTV CP77 ortholog has a large deletion at the 5’ end that results in an early frameshift. On the other hand, MPXV, TATV, AKMV, SKPV, and VPXV grew in both the parental and ΔSAMD9L CHO cells with nearly the same efficiency, indicating that chSAMD9L did not pose significant restriction for their replication. These OPXV species all encode a full-length CP77 ortholog, with the two North American OPXVs (SKPV and VPXV) having the most divergent ortholog of ∼70% aa identity to CPXV CP77. RCNV, another North American OPXV, also grew in both the parental and ΔSAMD9L CHO cells, but the yield was more than 10-fold less in the parental cells at both 24 and 48 h post infection (Fig. 7), indicating that chSAMD9L reduced but did not block RCNV growth in CHO cells. Interestingly, a gene fusion event in RCNV genome resulted in an ORF with the N-terminal 406 aa of the CP77 ortholog and the 246 aa chemokine binding protein. Altogether, the data show that the ability of OPXV species to antagonize chSAMD9L and grow in CHO cells correlates with their coding of a CP77 ortholog with at least the N-terminal 406 aa.

**Table 1.** OPXV CP77 orthologs orthologs comparison to CPXV BR CP77.

| OPXV species | CP77 |  | Antagonize chSAMD9L |
| --- | --- | --- | --- |
|  | length (aa) | identity (%) |  |
| **CPXV** | 661-674 | 91-100 | Yes |
| **MPXV** | 659-660 | 90-91 | Yes |
| **VACV** | small deletions, early frameshift |  | No |
| **VARV** | >600 nt deletion, 355-490 | 86-88 | ND |
| **ECTV** | 504 nt deletion, early frameshift |  | No |
| **CMLV** | large deletion (~155 nt remaining) |  | No |
| **TATV** | 661 | 92 | Yes |
| **AKMV** | 672 | 79 | Yes |
| **SKPV** | 633 | 69 | Yes |
| **VPXV** | 613 | 69 | Yes |
| **RNCV** | gene fusion*, 406 | 66 | partial |

\*Strain MD1964-85A. 1-406 of CP77 ortholog fused in frame to the N-terminus of the full-length, 246-aa chemokine binding protein. Total protein length is 655-aa.

**Figure 7.**
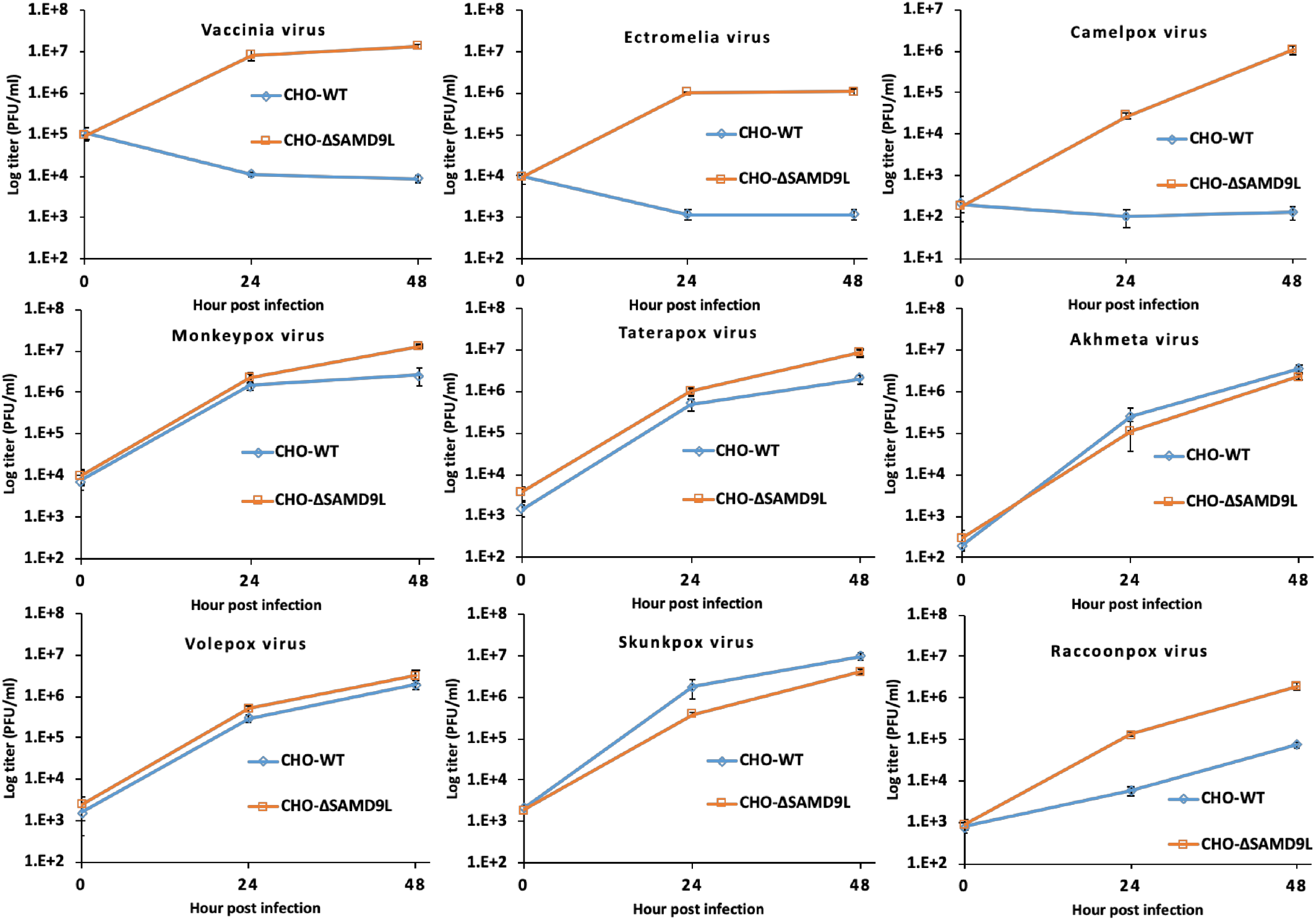
Orthopoxvirus host range in CHO cells corresponds to their CP77 gene status. The parental (CHO-WT) and ΔSAMD9L CHO cells (CHO-ΔSAMD9L) were infected with the indicated orthopoxvirus species. Viral titers at 0, 24 and 48 hpi were measured by plaque assay on VERO cells.

## DISCUSSION

Many infectious diseases that result in high morbidity and mortality in humans are zoonoses. VARV, an exclusive human pathogen, is believed to have evolved from an African rodent-borne virus (41), before it spread around the world and became mankind’s deadliest killer. For many viruses, divergence in viral entry receptors in different host species poses a major hurdle for cross-species transmission (42). Poxviruses, however, can enter nearly any animal cell (43). Why many poxviruses show strict host species specificity are less clear, but some host antiviral factors such as PKR and SAMD9L have shown some species-specific difference in susceptibility to poxvirus inhibitors (25, 44, 45). In this study, we revealed a species-specific difference in SAMD9L as the cause for the restriction of several OPXVs (VACV, ECTV, CMLV) in a rodent cell, suggesting that divergence in SAMD9L (and perhaps SAMD9) in rodent species presents a major barrier for cross-species poxvirus infection. Furthermore, we identified CP77 as the third OPXV SAMD9L inhibitor with a unique species specificity, demonstrating the sophistication brought forth by OPXVs to antagonize SAMD9&L.

OPXVs have broad host range in tissue culture cells, but CHO cells, while permissive for CPXV (16), are nonpermissive for VACV (14) and ECTV (46). What host factor causes the restriction of some OPXV species in CHO cells had been enigmatic since the initial discovery more than 50 years ago (14). In this study, we solved this mystery by identifying chSAMD9L as the host restriction factor in CHO cells. We presented two complementary lines of evidence: 1) CRISPR-Cas9 KO of chSAMD9L from CHO cells completely removed the host restriction for VACV, ECTV and CMLV; 2) ectopic expression of chSAMD9L in a human cell line recapitulated the poxvirus restriction property of CHO cells. We also provided a molecular explanation why chSAMD9L could restrict some OPXV species, namely its resistance to binding by both K1 and C7, the two SAMD9&L inhibitors in these OPXV species. Interestingly, while SAMD9 is the constitutive poxvirus restriction factor in many human cells (23-25), chSAMD9 appears not to contribute to poxvirus restriction in CHO cells. KO of chSAMD9L alone is sufficient for abolishing the restriction of a panel of VACVs, most of which did not contain K1, the only protein that was shown to bind chSAMD9. Moreover, KO of chSAMD9 had no effect on poxvirus restriction in CHO cells. CHO cells thus resemble mouse cells more than human cells in that SAMD9L is the constitutive restriction factor (25). It is unclear why chSAMD9 does not restrict poxviruses in CHO cells. We speculate that either chSAMD9 has lost the antiviral function due to some lineage-specific mutations or chSAMD9 expression level in CHO cells is not sufficiently high. The former scenario would be analogous to the fate of mouse SAMD9, which had suffered a mouse lineage-specific gene loss (47). The latter scenario would be similar to human SAMD9L, which has to be induced to a high expression level by IFN to impose restriction on vK1^−^C7^−^ (25). However, our preliminary experiments of treating ΔSAMD9L CHO cells with mouse or universal type-1 IFN did not result in restriction of vK1^−^C7^−^.

CPXV BR CP77 was found to rescue VACV replication in CHO cells more than 40 years ago (16). Since then, a number of molecular functions have been ascribed to CP77, including the binding to host HMG20A, NF-?B subunit p65 and the SCF complex (17, 40). However, the molecular mechanism underlying the host range function of CP77 remained elusive. We showed in this study that CP77 co-immunoprecipitated chSAMD9L and rescued VACV replication in a human cell line that was induced to express chSAMD9L, demonstrating that CP77 is a SAMD9L inhibitor. Only the N-terminal 382 aa containing the first seven ankyrin repeats was essential for chSAMD9L binding, while deletion of ankyrin repeat 5 abolished the binding. Correspondingly, a CP77 mutant with only the first seven ankyrin repeats but not the one with deletion of ankyrin repeat 5 could rescue VACV replication in CHO cells, indicating that the host range function of CP77 relies on its binding with chSAMD9L. This idea was further supported by comparing the growth of nearly all OPXV species in the parental and ΔSAMD9L CHO cells. OPXV species that encode a full-length CP77 ortholog (MPXV, TATV, AKMV, SKPV and VPXV) can replicate in both the parental and ΔSAMD9L CHO cells, while OPXV species that have lost CP77 gene (VACV, ECTV, and CMLV) can only replicate in ΔSAMD9L CHO cells. Interestingly, RNCV, which encodes a fusion protein that contains only the N-terminal 406 aa of CP77 ortholog (48), can also replicate in CHO cells, albeit with reduced efficiency compared to that in ΔSAMD9L CHO cells. The RNCV genome is closely related to that of the other two North American OPXVs, with the largest difference a 25 kbp deletion in the left terminal region of RCNV that removed 12 complete genes and created an in-frame gene fusion of CP77 ortholog and the chemokine binding protein (48). The difference between the three North American OPXVs in terms of their replication in CHO cells correlates with our observation that the N-terminal seven ankyrin repeats was less effective than the full-length CP77 in inhibiting chSAMD9L. VARV encodes a CP77 ortholog varying from 355 to 490 aa in length, so VARV is predicted to be able to partially inhibit chSAMD9L.

Recently, we and others have established the importance of SAMD9&L as host restriction factors against poxviruses at the cellular and organismal level (23-25). The identification of CP77 as yet another SAMD9L inhibitor underscores the critical role of SAMD9&L in host defense against poxviruses and the elaborate lengths OPXVs went to evade SAMD9&L. K1 and C7 were previously shown to function equivalently at inhibiting human and mouse SAMD9&L (25). In this study, however, we uncovered differences between K1/C7/CP77 in their targeting mechanism and binding specificity for SAMD9&L. While a C7 ortholog was shown to target the N-terminus of SAMD9 (39), both K1 and CP77 target an internal region containing the predicated NTPase and TPR domains. While any one of K1/C7/CP77 can bind mouse SAMD9L, only K1 can bind chSAMD9, and only CP77 can bind chSAMD9L. SAMD9L from mouse and Chinese hamster shares ∼80% aa sequence identity. We speculate that a need for overcoming SAMD9&L sequence divergence in different rodent species may have driven OPXVs in evolving three different inhibitors targeting different regions of SAMD9&L. The loss of K1 or/and CP77 from OPXV species with a narrow host-range (VARV, CMLV) and the maintenance of all three (K1/C7/CP77) in species presumably endemic in wild rodents (CPXV, MPXV and North American OPXV) also suggest that multiple SAMD9&L inhibitors are needed specifically for overcoming diverse SAMD9&L in rodents.

## ACKNOWLEDGMENTS

This work was supported by a grant from NIAID to Y. Xiang (AI079217). We thank Nadia Gallardo-Romero for providing the OPXVs.

The findings and conclusions in this report are those of the authors and do not necessarily represent the official position of the Centers for Disease Control and Prevention.

